# Reconstitution of the uterine immune milieu after transplantation

**DOI:** 10.1101/2024.03.04.583277

**Authors:** Benedikt Strunz, Martin A. Ivarsson, Dan Sun, Christoph Ziegenhain, Ylva Crona-Guterstam, Martin Solders, Andreas Björklund, Nicole Marquardt, Helen Kaipe, Angelique Flöter-Rådestad, Sebastian Gidlöf, Mats Brännström, Niklas K. Björkström

## Abstract

Maintenance of tissue-specific immunity is important for immunological fitness, but its establishment have been difficult to assess in humans. Here, we investigated reconstitution of the human uterine immune system by studying women undergoing uterus solid organ transplantation (UTX) or hematopoietic stem cell transplantation (HSCT). Through single-cell identification based on SNPs and disparate HLA expression using single-cell RNA sequencing or high-parameter flow cytometry, donor vs recipient cell origin was determined, and features of these cells were studied. A full uterine immune cell reconstitution occurred after both UTX and HSCT, both at transcriptomic and phenotypic level. This occurred despite tacrolimus-induced calcineurin-mediated NFAT pathway inhibition, which affected de novo induction of tissue-residency features *in vitro*. Intriguingly, after HSCT, immune cells of male origin could reconstitute the uterine immune milieu. Collectively, our results proved insights into tissue immune system persistence and reconstitution capabilities in an organ undergoing continuous regeneration.

## Introduction

Human organs are populated by resident myeloid and lymphoid immune cells with a composition that differs between tissues, likely due to distinct anatomical factors and environmental cues (*1–3*). Compared to the dogma from murine parabiosis models where tissue residency is long-lasting (*4*), solid organ transplantation studies in humans have revealed an alternative picture with a greater degree of immune cell reconstitution over time (*5–8*). However, the reconstitution rate appears to be organ and cell-type specific with more rapid reconstitution of liver myeloid and lymphoid cells (*7*, *8*) whilst intraepithelial T cells remain donor-derived for longer periods of time after lung and intestinal transplantation (*5*, *6*, *9*) as well as of host origin in skin after hematopoietic stem cell transplantation (HSCT) (*10*). Yet, for other organs, if and under what circumstances immune reconstitution occurs remains elusive.

The human uterus contains a distinct immune cell composition, undergoes continuous regeneration throughout each menstrual period, and is further modified in pregnancy (*11*, *12*). For instance, uterine NK (uNK) cells, resident in this tissue, exhibit a continuous differentiation and expansion process in response to endometrial regeneration (*12*). This is coupled to progesterone influencing local stromal cells to produce and present IL-15 (*13*, *14*). Regarding the cyclic contribution of sex hormones to local immune homeostasis (*12*, *14–16*), little is known about cellular reconstitution. Three non-mutually exclusive hypotheses can be put forward: i) immune cell progenitors exist in the basal layer of the endometrium that remain throughout menstrual cycles, ii) bone marrow-derived progenitor cells home to the uterus at the start of each menstrual cycle and differentiate locally, or iii) peripheral blood-derived mature immune cells enter the tissue and differentiate into resident cells *in situ* (*17*).

To address uterine immune system reconstitution in humans, we studied two unique cohorts: women undergoing uterus transplantation (UTX) because of Mayer-Rokitansky-Küster-Hauser (MRKH) Syndrome (rare congenital reproductive disorder defined by a nonexistent uterus) (*18*, *19*) and women with recommenced menstruation after HSCT. We show that new immune cells enter the tissue and fully reconstitute the local microenvironment with new tissue resident cells that are transcriptionally and phenotypically similar to conventional uterine immune cells. This occurred despite an inhibited calcineurin – NFAT pathway and even immune cells of male origin could reconstitute the niche and respond to cyclic sex hormone signals. Our results demonstrate principles for uterine immune system reconstitution in humans.

## Results

### Assessment of the uterine immune milieu after UTX or HSCT

In this study, two unique cohorts of transplantation patients were recruited to study uterine immune cell reconstitution patterns. The first cohort comprised of six women having undergone uterus transplantation (UTX) and the second cohort of four women with retained ovarian function after allogeneic HSCT (**Fig. 1**). All UTX patients had undergone successful pregnancies after transplantation and either decidua tissue, obtained at the time of delivery by cesarian section (n=3), or endometrial tissue obtained at hysterectomy of the transplant after successful pregnancy (n=3) was studied (Table S1). Three out of four HSCT patients had successfully become pregnant and given birth before menstrual blood samples were obtained for the study (Table S2). As control material, decidua, endometrium, and menstrual blood samples were also obtained from unrelated healthy controls. UTX allowed us to study the uterine immune composition of a donor uterus within the recipient’s environment, whilst HSCT permitted the assessment of the cognate uterus when hematopoietic stem cells had been transplanted. To determine the uterine immune composition after transplantation, we performed high-dimensional flow cytometry of immune cells in endometrium, decidua, and menstrual blood of UTX and HSCT samples and compared the results to control cohorts (**Fig. 2A-E**). After UTX and HSCT, major immune cell linages including myeloid cells, T cells, NK cells, and B cells could readily be identified, and levels were not significantly different to those seen in controls (**Fig. 2A-B**). UMAP analysis comparing contributions of immune cells from UTX/HSCT and healthy controls further revealed that all clusters were populated by cells from both sources and that only minor differences were present compared to controls (**Fig. 2C**).

**Fig 1.**
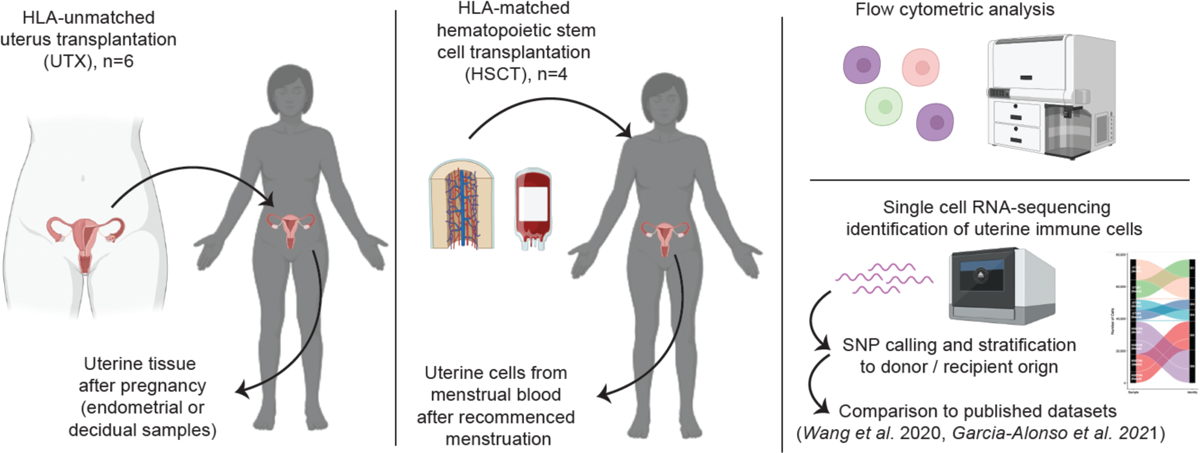
Study overview. In this study two cohorts of transplantation patients were included: patients with solid organ uterus transplantation (UTX, n=6) and patients after hematopoietic stem cell transplantation (HSCT, n=4). Menstrual blood and uterine tissue samples were analyzed using flow cytometry and single-cell RNA sequencing to determine the uterine immune composition as well as donor or recipient origin.

**Fig. 2.**
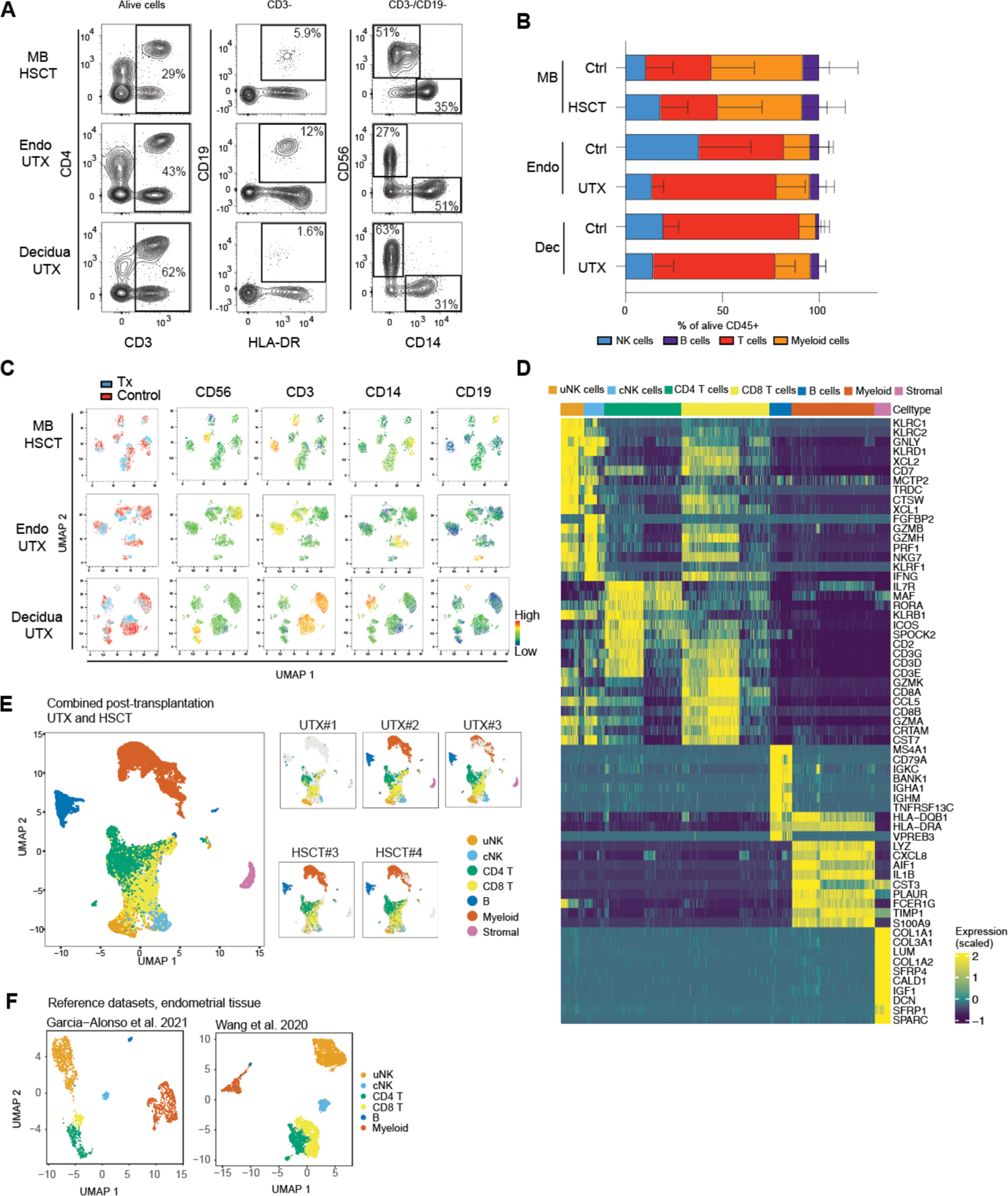
The composition of the uterine immune milieu after transplantation is comparable to healthy control samples. (**A-C**) Flow cytometric assessment of immune cells in the uterus of UTX and HSCT patients or respective controls, studied were menstrual blood (MB) in HSCT, endometrium (Endo), and decidua (Dec) in UTX patients. (**A**) Representative flow cytometry plots of the major immune cell subsets and (**B**) summary plot depicting the frequency of alive CD45^+^ cells as mean with standard deviation (n for controls/patients: MB n=5/4, Endo n=6/3 and Dec n=5/3). (**C**) UMAP analysis comparing the phenotype of uterine immune cells in transplantation patients and controls. (**D**) Gene-expression profile identifying the major cell types in combined analysis of all patients (uNK = uterine NK cells, cNK= circulating NK cells). (**E**) UMAP analysis of combined and individual samples and (**F**) of two reference datasets of uterine immune cells from *Garcia-Alonso et al.* (Nature Genetics, 2021) and *Wang et al.* (Nature Medicine, 2020).

Next, we performed single-cell RNA sequencing (scRNAseq) of endometrial and menstrual blood cells of UTX and HSCT, respectively (**Fig. 2D-F**). In line with the flow cytometric analysis, immune cells from UTX and HSCT clustered together and delineated the major immune cell subsets expected in the uterus (**Fig. 2E**). scRNAseq of UTX#1 was performed on FACS-sorted T cells and NK cells and thus only those were recovered. UTX#2 and #3, in which live CD45+ cells from the endometrium were subjected to scRNAseq, also contained other immune cells as well as a small fraction of stromal cells. Menstrual blood HSCT samples had fewer stromal cells (**Fig. 2E**). As a control we applied our annotation of the identified immune cell subsets to two publicly available datasets generated from endometrial tissue (*20, 21*), which resulted in comparable clustering of the uterine immune cells (**Fig. 2F**). In summary, all major immune cell subsets are present in non-pregnant and pregnant uterus after UTX and HSCT at levels comparable to healthy controls.

### Identification of donor and recipient origins of uterine immune cells after transplantation

Next, we leveraged single-nucleotide polymorphisms (SNPs) unique either to donor or recipient to determine origin of cells obtained from the scRNAseq analysis (**Fig. 3A**). For UTX, SNPs from peripheral blood cells of the uterus-donor were used as a reference. Interestingly, the vast majority of uterine immune cells in UTX samples were not of donor but of recipient origin (**Fig. 3B-C**) suggesting that the uterine microenvironment had been reconstituted by recipient-derived cells. Recipient-derived cells also dominated all discrete immune cell subsets (**Fig. 3D**) whilst stromal cells remained of donor origin (**fig. S1)**. In case of HSCT, all uterine immune cells after HSCT were of donor origin (**Fig. 3B-C, and E)**. Thus, similar to UTX, also after HSCT the uterine microenvironment is reconstituted by de novo-generated immune cells.

**Fig. 3.**
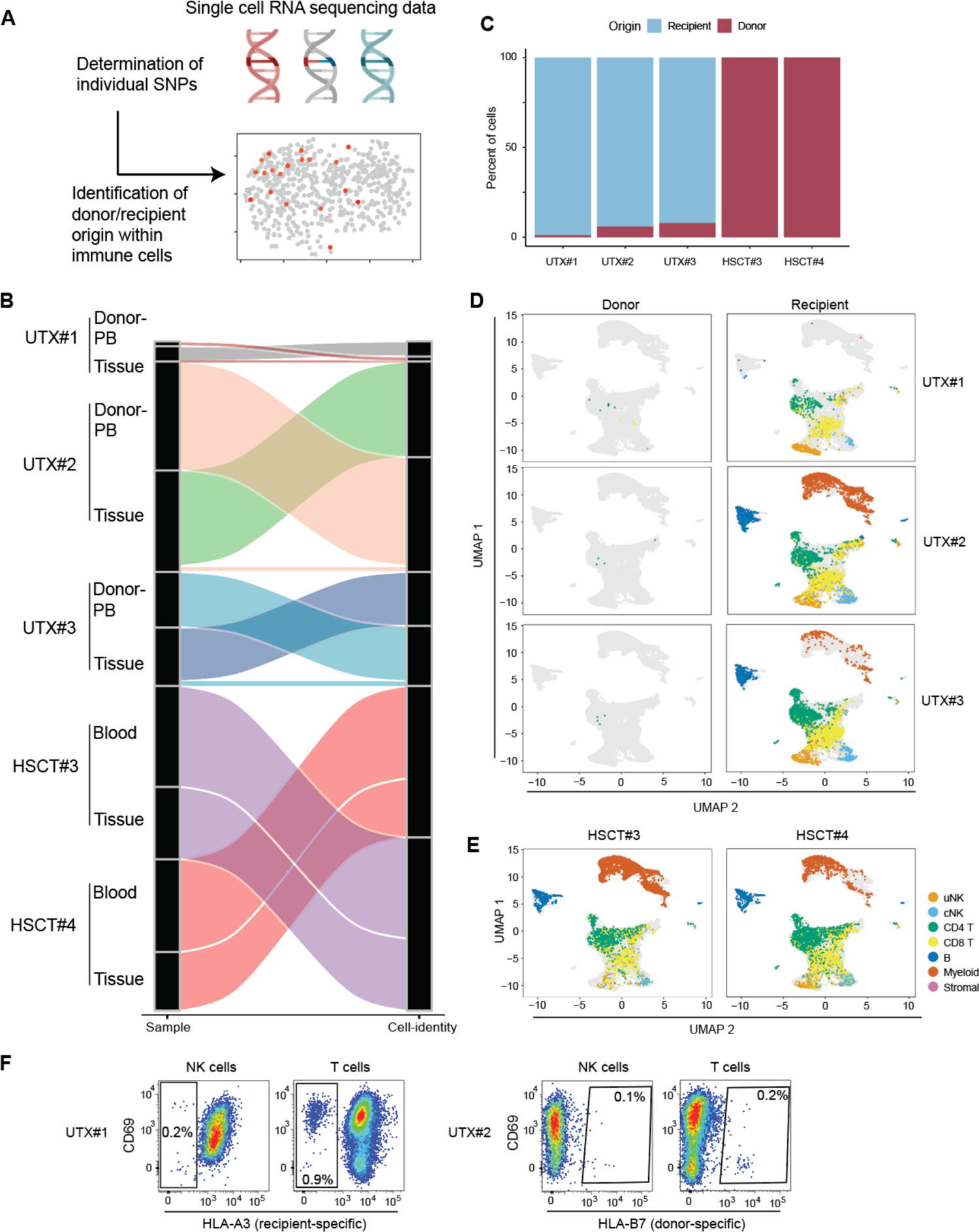
Identification of donor- and recipient-identity of uterine immune cells after transplantation. (**A**) Schematic displaying the workflow for allocating cells to unique individuals. (**B**) Summary of the identity of uterine immune cells in UTX and HSCT. (**C**) Summary showing relative frequency of cells stratified for donor/recipient identity. (**D** and **E**) UMAP depiction of uterine immune cells stratified for donor/recipient origin in UTX (**D**) and HSCT (**E**, all cells of donor origin). (**F**) Flow cytometry analysis of UTX#1 and UTX#2 live uterine CD45+ immune cells. The uterus was transplanted with HLA-mismatch and either recipient-specific (left hand side, UTX#1) or donor-specific (right hand side, UTX#2) HLA could be stained.

The results from the UTX samples were also validated using flow cytometry and staining for donor/recipient-specific HLA (**Fig. 3F**). Sampling of the UTX patients was performed at least one year after the transplantation (Table S1) but small fractions of donor-derived immune cells could still be identified both using scRNAseq and flow cytometry (**Fig. 3B-D, F**). These cells were primarily of NK cell and T cell origin and no donor-derived myeloid or B cells could be identified (**Fig. 3D and F**). No residual immune cells of recipient origin could be detected in the uterus after HSCT.

In summary, whilst small numbers of donor cells can persist in the uterus up to several years after UTX, most of the uterine immune cell compartment gets replenished by recipient immune cells. After HSCT, the uterus gets solely reconstituted by donor-derived immune cells.

### Reconstitution of tissue-resident uterine immune cells after transplantation

Since most immune cells in the uterine microenvironment had been replaced following UTX or HSCT, we next investigated these reconstituted cells in more detail. We focused on NK cells, CD4 and CD8 T cells and first performed separate sub clustering of the scRNAseq data for each cell population (**Fig. 4A, fig. S2**). Two clusters of NK cells could be identified annotated as either uterine (uNK) or conventional NK cells (cNK). uNK cells showed high expression of transcripts for the tissue-residency markers CD49a (*Itga1*) and CD103 (*Itgae*) (**Fig. 4B)** and enriched for a canonical tissue-residency gene signature (**Fig. 4C**). Three clusters of CD4 T cells could be identified, regulatory T cells (Tregs), tissue-resident memory cells (CD4 TRM), and a combined naïve and central memory cluster (CD4 Naïve/CM) (**Fig. 4A**). Within CD8 T cells, mucosal-associated invariant T (MAIT) cells, two clusters of CD8 TRMs, as well as one CD8 effector memory (TEM) and one naïve/CM cluster were revealed. Similar to NK cells, CD4 and CD8 TRM-clusters had high transcript levels of tissue-resident markers (CD49a and CD69) and were enriched for a canonical TRM gene signature (**Fig. 4B and D, fig. S3A)**. Interestingly, for T cells, the *de novo* reconstituted TRMs enriched for this gene signature to a similar degree as the few remaining donor derived cells **(Fig. 4D**).

**Fig. 4.**
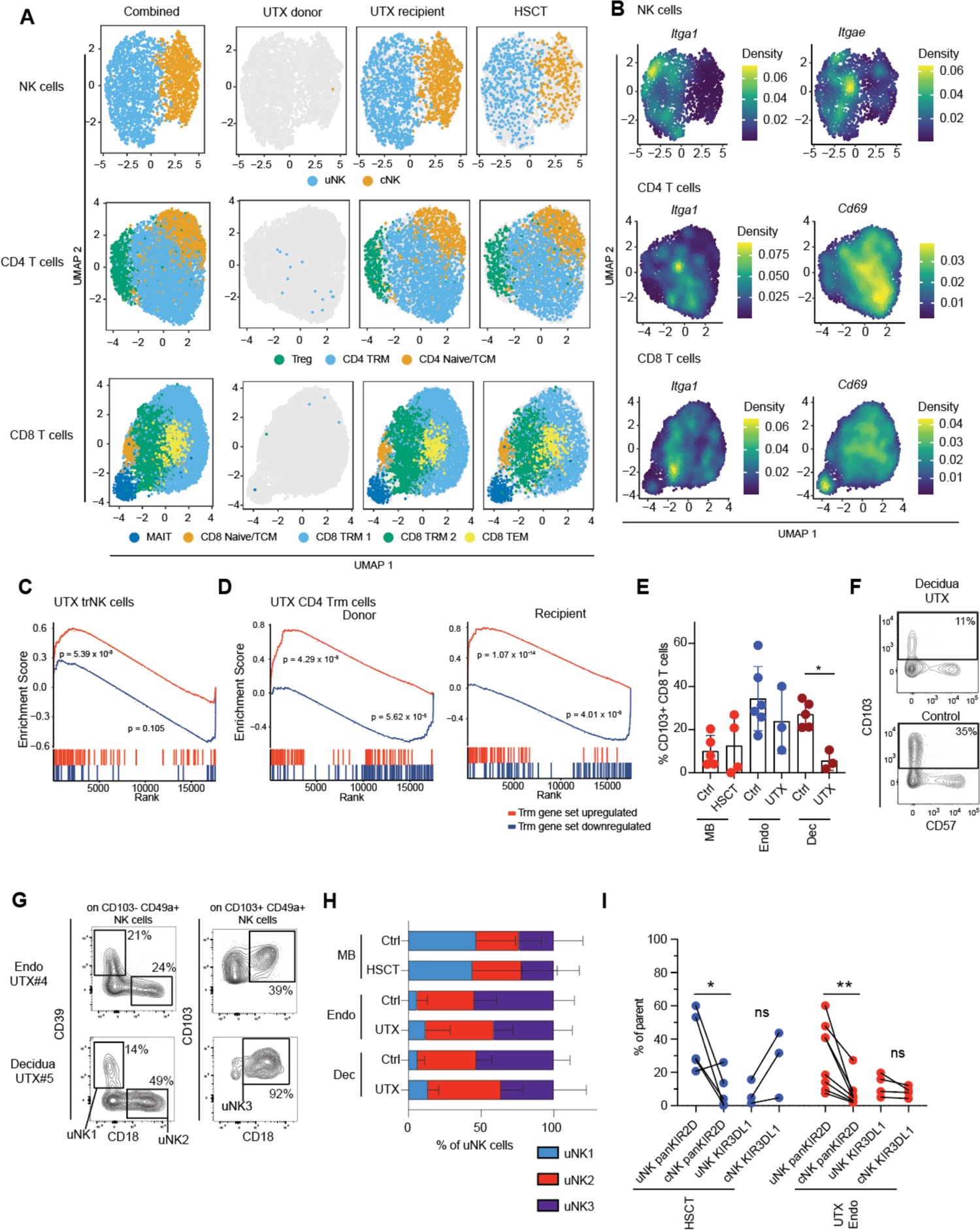
Reconstitution of tissue-resident uterine immune cells after transplantation. (**A**) UMAP analysis of integrated 10x data showing NK, CD4, and CD8 T cells stratified for donor/recipient identity and circulating (cNK, CD4 Naïve/TCM, CD8 Naïve/TCM, CD8 EM) or tissue-resident/uterine (uNK, CD4 TRM, CD8 TRM) gene expression profile. (**B**) Expression of indicated tissue-residency related genes overlayed on the cell types displayed in (**A**). (**C** and **D**) Gene set enrichment analysis for tissue-residency related gene sets in donor/recipient-originating NK cells (**C**) as well as T cells (**D**). (**E** and **F**) Analysis of CD103 expression on CD8 T cells in the indicated groups, (**E**) summary and (**F**) representative flow cytometry plots. (**G**-**I**) Flow cytometric analysis of uterine NK (uNK) cell phenotype. (**G**) Representative gating on uNK cell subsets 1-3 in endometrial and decidual UTX samples. (**H**) Summary of uNK subset frequency in the indicated cohorts/controls, displayed is relative frequency in relation to total uNK1-3. (**I**) NK cell subsets expressing any KIR2D receptors (KIR2DL1/S1, KIR2DL2/L3/S2 and KIR2DS4, n=4 for HSCT and n=8 for UTX Endo) vs. KIR3DL1 (n=4 for HSCT and n=3 for UTX endo), stratified for uterine or circulating cell origin. Tested for statistically significant differences with either paired t-test or Wilcoxon test, * indicates p<0.05 and ** p<0.01.

Within the uterus, uNK cells and macrophages are frequent compared to other organs and to blood (*11, 22*). Thus, we also investigated receptor-ligand interaction between uNK cells and macrophages using CellphoneDB as well as cytokine signaling pathways with Cytosig (**fig. S3B**). Results from UTX and HSCT were compared to data from two public datasets. In line with our observations regarding composition and tissue residency profiles, both cellular interactions and deducted cytokine signaling pathways were at an overall level comparable in UTX and HSCT to that of healthy controls, yet minor differences could be observed (**fig. S3B**). Corroborating the scRNAseq data, presence of CD103-expressing T cells could be validated using flow cytometry, albeit with a lower fraction of such cells in UTX decidua compared to controls (**Fig. 4E, F**). uNK cells can be subdivided into less and more differentiated uNK cells based on CD39, CD18, and CD103 expression (referred to as uNK1, 2, and 3, (*11, 12*)). We could identify all three uNK cell subsets in menstrual blood, endometrium, and decidua after transplantation (**Fig. 4G an H)** and, despite differences between menstrual blood and uterine tissue, fractions of uNK cell subsets were comparable to the respective controls (**Fig. 4H).** Another feature of uNK cells (here, predominantly endometrial NK cells in the non-pregnant uterus) is the skewed KIR-profile marked by more frequent KIR2D expression on the expense of KIR3D-receptors (*22–24*). Indeed, also skewing of the KIR2D-profile was recapitulated on uNK cells after transplantation, while reduced expression of KIR3D-receptors could be observed mainly in HSCT samples (**Fig. 4I)**. In summary, tissue-resident T cells and uNK cells can transcriptionally and phenotypically be *de novo* reconstituted in the uterus after UTX and HSCT.

### Tissue-resident NK cells are formed despite calcineurin – NFAT signaling inhibition

Following UTX, the immunosuppressant tacrolimus was used to avoid rejection (Table S1). Tacrolimus inhibits the calcineurin – NFAT signaling pathway (*25*). Interestingly, in UTX-patients uNK cells displayed a significantly downregulated NFAT-signature as compared to cNK cells (**Fig. 5A, fig. S4A)**. However, this was not the case for HSCT-patients or healthy control datasets (**fig. S4A and B)**. Similarly, the NFAT-signature was significantly enhanced among CD4 TRM cells from controls and HSCT whilst this enhancement was absent in CD4 TRM from UTX-patients (**Fig. 5B, fig. S4B**). Of note, no alteration in CD8 TRM was observed (fig**. S4C**). Collectively, this suggests that tacrolimus had a greater effect on tissue-resident compared to circulating lymphocytes and more on NK cells than T cells. Yet, tissue-resident lymphocytes could replenish the uterine niche (**Fig. 4**). This would suggest that NFAT-signaling is of lesser importance for establishment of tissue-resident cells in the uterus. To functionally address this, we cultured purified peripheral blood NK cells with IL-15 and TGFβ, a combination known to induce a tissue-residency/uterine phenotype of NK cells (*26*), with or without tacrolimus. Of interest, proliferation and induction of tissue-residency markers associated with uNK cells was significantly affected by tacrolimus treatment (**Fig. 5C-E, fig. S5A)**. In detail, tacrolimus treatment lead to a significantly reduced induction of CD49a, CD69, CD103, CD39 and CD9 **(Fig. 5D and E, fig. S5A)**. Thus, although tacrolimus dampens calcineurin – NFAT signaling in tissue-resident lymphocytes and *in vitro* tacrolimus treatment formation of tissue residency (**Fig. 5**), *de novo* generation of tissue-resident immune cells still occurred over time *in vivo* (**Fig. 4**).

**Fig. 5.**
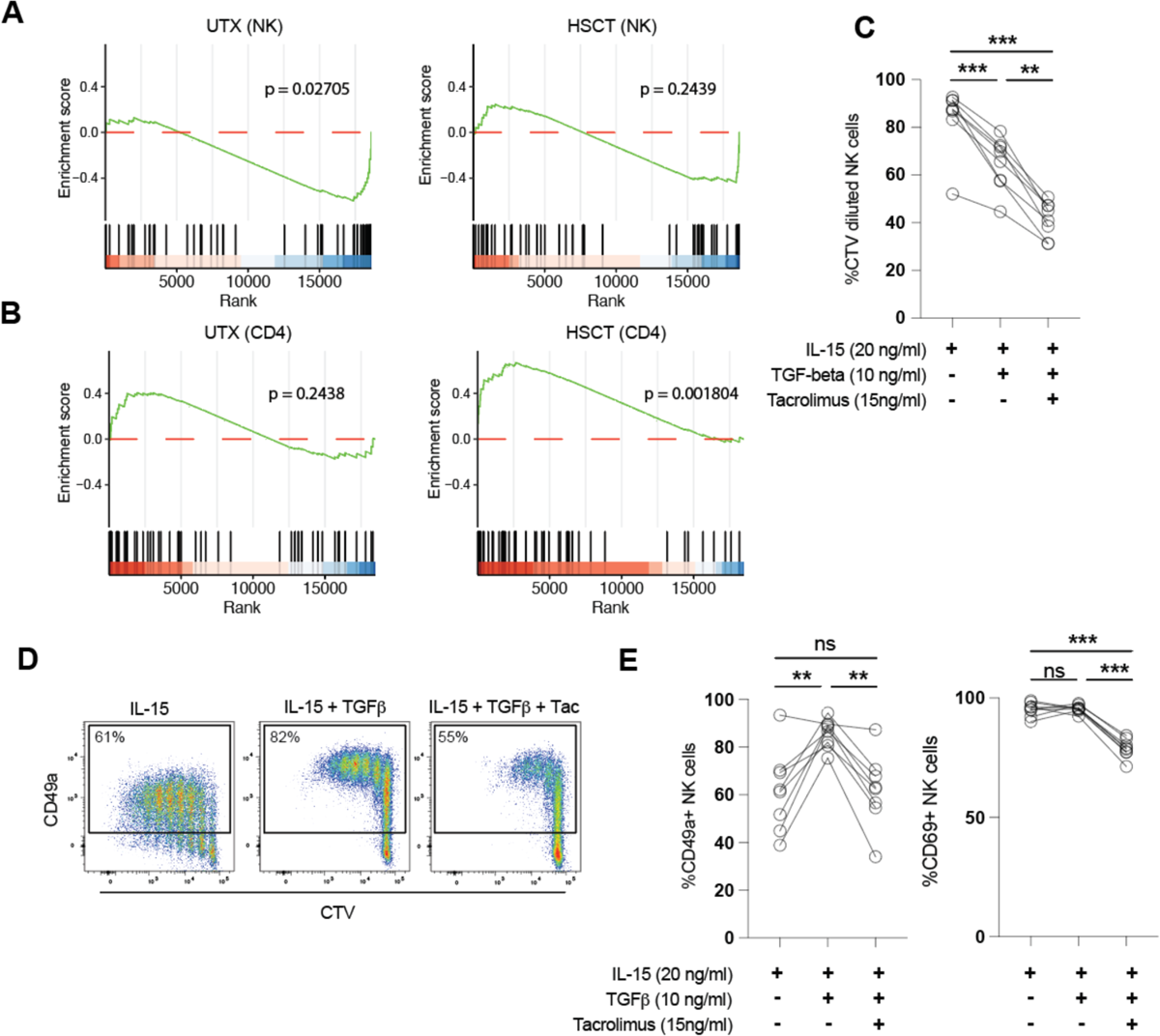
Calcineurin-inhibition predominantly influences tissue resident immune cells. (**A-B**) Gene set enrichment analysis of NFAT signaling pathway in NK cells (**A**) and CD4 T cells (**B**). (**C** - **E**) Effect of Tacrolimus treatment on IL-15/TGF-beta mediated proliferation (**C**) or CD49a and CD69 upregulation (**D-E**) on NK cells. Tested for statistically significant differences with ANOVA followed by Tukey’s test for multiple comparisons, ** indicates p<0.01, *** p<0.001.

### An immune system of male origin can form tissue-resident uterine immune cells

In one HSCT patient (HSCT#4), the hematopoietic stem cell graft came from a male donor. Hence, this gave us the unique opportunity to assess if *bona fide* uNK cells and other tissue-resident lymphocytes could reconstituted the uterine microenvironment despite being of male origin. The immune cell subset composition based on scRNAseq analysis did not differ in this HSCT-patient (**Fig. 6A**) compared to other transplantation donors (**Fig. 4A**). Next, we looked at expression of transcripts located on the Y-chromosome in uterine immune cell subsets. We observed expression of Y-chromosome transcripts (*Ddx3y, Eif1ay, Uty*) both in tissue-resident NK and T cells as well as circulating counterparts (**Fig. 6A**). uNK cells of male origin also enriched strongly for a canonical tissue-residency immune signature (**Fig. 6B**). We have previously shown that uNK cells differentiate in response to progesterone and IL-15 during each menstrual cycle and acquire features not present in other organs (*12*). To determine if uNK cells of male origin also could undergo such cyclic differentiation, presence of uNK1-3 subsets was determined in HSCT#4. Indeed, these subsets of uNK cells were present at comparable levels as in control menstrual blood samples (**Fig. 6C**). Finally, we could also show that formation of tissue residency markers and uNK cell markers occurred at similar efficiency *in vitro* in response to IL-15 and TGBβ in peripheral blood NK cells when stratifying for the genetic sex of the donor (**fig. S5B)**. Thus, beyond that the circulating immune system could reconstitute the uterine niche following UTX, we here also show that a bone marrow of male origin can achieve this at a similar level.

**Fig. 6.**
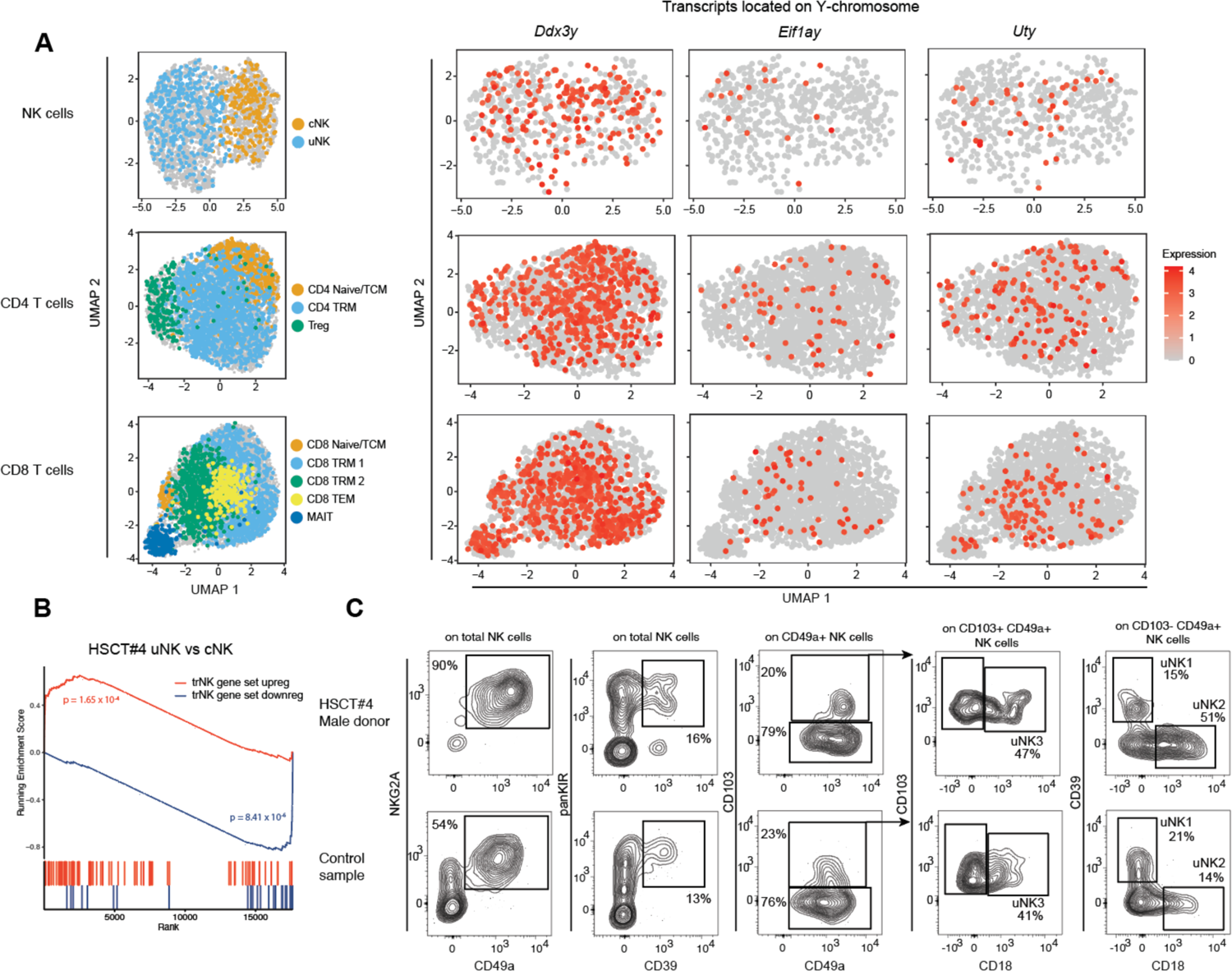
Formation of uterine immune cells regardless of genetic sex. Detailed analysis of HSCT#4 uterine immune cells after transplantation of stem cells from a male donor. (**A**) Depiction of UMAP analysis with uterine immune cells of HSCT#4 (left hand side, cells from all other samples gray in background) and overlayed expression of the indicated Y-chromosome-located transcripts in donor HSCT-4. (**B**) Enrichment of tissue-residency gene sets in uterine and circulating NK cells (uNK and cNK, respectively) in donor HSCT#4. (**C**) Flow cytometric analysis displaying uNK cell subsets in HSCT-4 and control uterine NK cells.

## Discussion

The uterine immune environment is unique in its composition and function, but how it is formed and whether it can be reconstituted is poorly understood. Here, we have addressed this by studying two cohorts of patients having undergone either solid organ uterus transplantation or hematopoietic stem cell transplantation. This allowed us to assess the possible reconstitution of the uterine immune milieu in settings where either the organ itself or the immune system had been replaced, and to query if de novo-generated immune cells could be detected in a post-transplantation setting.

Similar to other organs, the uterus displays a unique immune cell composition with a high frequency of specialized NK cells (*1, 11, 12, 22*). We show that the uterine immune niche is reconstituted after both transplantation settings. In more detail, the reconstitution occurred with immune cells present at similar frequencies as in non-transplanted controls. The reconstitution occurred both at the phenotypic and transcriptomic and phenotypic level, including tissue resident T cells and uNK cells present in the organ. This was also the case both in the non-pregnant (endometrium and menstrual blood samples) as well as in the pregnant (decidua) setting. This shows the importance of the local organ niches in tailoring tissue immune composition and suggests that this not only happens during initial immune development early in life but rather continuously throughout life.

Most cells (>95%) across immune linages were reconstituted after UTX or HSCT. These data further extend and corroborate previous more limited reports in few individuals showing that MAIT cells and KIR^+^CD39^+^ uNK cells are of recipient-origin after UTX (*12, 27*) and that CD45+ donor-derived cells are present in the endometrium after HSCT (*28*). Yet, few donor-derived T cells enriching for a TRM gene signature could be detected long after uterus transplantation. In contrast, very few, if any, uNK cells were donor-derived after UTX and all immune cells had been replaced after HSCT. Longevity and replenishment rates have previously been characterized more in detail for T cells in different transplantation settings including liver (*8*), lung (*5*), and intestine (*6, 9*) after solid organ transplantation and skin (*10*) after HSCT. Longevity rates vary with up to half of T cells still being of the pre-transplant origin in skin, lung, and intestine years after transplantation while a more rapid replenishment occurs in the liver. This might be related to the characteristics of the spatial niches in these organs where a larger fraction of intraepithelial lymphocytes expressing CD103 are present in skin (epidermis), lung, and intestine as compared to liver (*29–31*). Indeed, the CD103 - E-cadherin interaction might actively retain cells in tissues (*32*). Although we show that smaller subsets of both T cells and uNK cells express CD103 in the uterus, levels found are typically more in line with the lower levels found on intrahepatic lymphocytes (*30*) and not as prominently expressed as in gut, lung, and skin (epidermis) (*5, 9, 10*). Notably, donor-derived, i.e. long-term resident T cells were still present in the uterus despite the organ having undergone numerous menstrual cycles as well as a pregnancy.

We here describe the reconstitution of the uterine immune milieu in a HSCT patient that had been transplanted with hematopoietic stem cells from a male donor. In scRNAseq analysis we observed expression of Y-chromosome transcripts and identified all uNK cell subsets at levels comparable to healthy controls. Endometrial regeneration is regulated by female sex hormones. Immune cells can either respond directly or indirectly to these. Estrogen-response elements are for instance broadly expressed on immune cells (*33*). Progesterone will also act on endometrial stromal cells inducing production of IL-15 that in turn drive uNK cell differentiation (*12, 13*). The presence of an intact uterine immune compartment of male origin suggests that Y-chromosome transcribed genes and absence of X-inactivation do not negatively regulate processes necessary for its formation.

The immunosuppressive drug tacrolimus was used after UTX to avoid rejection. It inhibits calcineurin – NFAT signaling (*25*) and blocks proliferation and effector functions in NK and T cells. Although Egr2, downstream of NFAT, has been shown to be necessary for development and maturation of NKT cells (*34*), mature PB NK cells were present after UTX. Similarly, tissue-resident T cells and NK cells could be *de novo* reconstituted despite calcineurin – NFAT signaling inhibition where tissue-resident cells had a suppressed NFAT pathway in UTX. This occurred even though *in vitro* experiments revealed a lower level of induction of a tissue-residency phenotype with upregulation of CD9, CD103 and CD49a when treated with tacrolimus. Indeed, Smad and NFAT signaling pathways have been shown to cooperate to induce CD103 (*35*). However, in the uterine context, the calcineurin – NFAT pathway appears redundant for formation of tissue residency.

Our study is limited by the relatively few subjects studied. However, the clinical settings of UTX as well as regained ovarian function after HSCT are incredibly infrequent. Yet, these clinical situations allow for a unique insight into uterine immunology and were studied in detail. Along the same lines, larger cohorts would have allowed to assess the contribution of uterus-infiltrating lymphocytes to rejection episodes observed after UTX. Furthermore, given the different materials studied here (menstrual blood, endometrium, and decidua) certain features might be missed in the respective group.

In conclusion, by studying rare cohorts of either uterus-transplanted or hematopoietic stem cell transplanted patients we report that the uterine immune environment can be reconstituted after transplantation. Furthermore, we could identify small populations of tissue-resident donor-derived T cells that persisted for years after transplantation, in a tissue undergoing continuous regeneration with each cycle, as well as pregnancy. Immune cells residing in the uterus are important determinants for pregnancy outcome and play fundamental roles in uterine disorders. Understanding the immune homeostasis of this niche is thus important for our understanding of uterine immunology and pregnancy.

## Materials and Methods

### Study design

This study included two patient cohorts (UTX and HSCT) and healthy controls. Oral and written informed consent was obtained before inclusion in the study, and the local Ethics Review Boards in Stockholm and Gothenburg approved the study. Samples from UTX patients were taken at the Sahlgrenska University Hospital, Gothenburg, Sweden, either from decidua when the child was born via planned *sectio*, or from the endometrium when the uterus was removed during planned hysterectomy (planned procedure as part of every uterus-transplantation (*18, 19*)). Endometrium or decidua was obtained from the inside of the uterus immediately after hysterectomy or at delivery. Samples were immediately put in cell culture medium and transported to the laboratory for analysis or cryopreservation for grouped analysis. Peripheral blood was also obtained from each donor and patient (except recipient blood for UTX1) and PBMCs were preserved alongside uterine material. UTX patients experienced in average 2 (0–4) rejection episodes in the first 12 months after transplantation and the median time from transplantation to embryo transfer was 18,5 months (12-45 months, Table 1). As controls to the UTX samples we obtained endometrial samples taken during routine hysterectomy (performed for non-malignant reasons such as fibroma) or samples taken from decidua following planned *sectio* (*36*), both collected at Karolinska University Hospital, Sweden. HSCT patients were sampled for menstrual and peripheral blood as previously reported (*23*). Regarding HSCT patients, three regained regular menses spontaneously and one induced menses via hormonal replacement therapy. The time from HSCT to spontaneous onset of menstruation ranged from 3 months to approximately 7 years (see table 2). One HSCT patient received immunosuppressive treatment with cyclosporin A during sampling and was included for flow cytometric analysis but not for single cell RNA sequencing. Control samples for the HSCT cohort were collected from healthy individuals with regular menses. Chronic disease and uterus anomalies were exclusion criteria. Menstrual blood from HSCT patients and controls were sampled at Karolinska University Hospital, Sweden.

### Sample processing

All samples were processed within 48 hours after collection. Peripheral blood samples were collected in EDTA or Heparin blood collection tubes, menstrual blood samples were collected within the first 24h of menstruation as previously reported (*23*). Endometrium UTX and decidua UTX samples were collected in RPMI (supplemented with 10% FCS, 2mM L-glutamin and Penicillin/Streptomycin), dissected in small pieces with scalpels before digestion with 0.25mg/mL collagenase II and 0.2mg/mL DNase I for a maximum of 30 minutes. Of note, cryopreserved control decidua samples were subjected only to mechanical dissociation as previously described (*36*). Subsequently, mononuclear cells from either tissue or blood samples were isolated with density gradient centrifugation. After washing cells were used either fresh for experimentation or cryopreserved.

### Flow cytometry, cell sorting and downstream analysis

Flow cytometry staining was performed as previously described (*12*). In short, 2*10^6 cells were resuspended in FACS Buffer (PBS +2% FCS +2mM EDTA), washed and stained for 20 minutes for surface markers (antibodies used for staining are listed in Table S3). Dead cells were identified using Fixable LIVE/DEAD Aqua or Yellow (ThermoFisher). In case of known discrepancies in HLA genotype between donor and recipient, specific HLA antibodies were used to discriminate donor and recipient lymphocytes. For intracellular staining, cells were fixed, permeabilized (ThermoFisher/FOXP3 Transcription staining buffer set, eBioscience) and stained intracellularly for 30 minutes. Data were acquired either on a BD LSR Fortessa XL or a BD Symphony with 5 lasers. Fluorescence-activated cell sorting (FACS) was performed on a Sony MA900. Downstream analysis, including UMAP analysis was performed in FlowJo version 10.

### Assessment of the effect of Tacrolimus or genetic sex on uterine NK cell generation

To study the effect of Tacrolimus on the generation of tissue residency profiles/uNK cell phenotypes, peripheral blood NK cells were isolated from PBMC obtained from healthy donors. Briefly, PBMC were thawed, and NK cells were isolated using magnetic negative isolation of NK cells (Miltenyi). Purified NK cells were labelled with Cell Trace Violet (CTV, Thermo Fisher) according to manufacturer’s instruction and plated at 1*10^6 cells/well. Cells were then stimulated for 5 days with indicated combinations of IL-15 (20ng/mL, RnD Biotechne), TGFβ (10ng/mL,RnD Biotechne), and Tacrolimus (Prograf, Astellas, 15ng/mL in PBS). For determination of the influence of genetic sex, PBMC with known sex of the donor were used. As readout, cells were stained and analyzed via flow cytometry as described above.

### Single-cell RNAseq analysis

In total three UTX and two HSCT samples were subjected to single cell sequencing. For the first UTX sample, we performed a targeted Fluorescence-activated single cell (FACS) sort to obtain certain leukocyte subsets either from donor (B cells) or recipient peripheral blood (monocyte), as well as T and NK cells from recipient tissue. Right before performing 10x genomics, cells were combined and subjected to 10x Genomics single cell sequencing via the SciLife single cell sequencing facility. The remaining two UTX and HSCT samples consisted of alive CD45+ sorted mononuclear cells either from tissue or from peripheral blood. cDNA and Gene Expression libraries were created according to the manufacturer’s protocol. 10x Genomics Gene Expression libraries were then converted to single-stranded circular DNA from 50 ng using the Universal Library Conversion Kit (App-A; MGI Tech) according to the manufacturer’s instructions. Libraries were pooled and 60 fmol of the pool was used to generate DNA nanoballs directly prior to sequencing on two FCL flow cells on the DNBSEQ G400RS platform (MGI Tech). We generated asymmetric paired-end reads (28-8-150 cycles). Library pools were also subjected to sequencing on a S2 flow cell using the NovaSeq6000 platform (Illumina) generating asymmetric paired-end reads (28-8-100 cycles).

### Single-cell RNA-seq data preprocessing, cluster identification and annotation

The demultiplexing of FASTQ files was carried out using deML v1.1.3. Post demultiplexing, the reads from each sample underwent quality filtering, alignment to the GRCh38 reference genome, quantification with GENCODE v35 gene annotations utilizing the zUMIs pipeline v2.9.4f. CellBender 0.1.0 was used to exclude empty droplets and mitigate background noise, while Solo 1.0 was applied to drop doublets. Genetic identities of recipient tissue and donor blood cells were delineated first by aggregating read counts on either reference or alternative alleles on approximately 7.4 million prevalent variants with an allele frequency surpassing 5% in the 1000 Genomes Project phase 3 release utilizing cellsnp-lite 1.2.0. Post genotyping, cells were sorted into a range of five to 10 genetic identities (--nDonor) using Vireo 0.4.2. Among these, eight was chosen as the optimal count given it represented the lowest number of identities while concurrently exhibiting the minimal number of genetic doublets. The genetic identities were then visualized utilizing the ggalluvial 0.12.3 package. The expression matrices from the each sample were log-normalized, and the normalized expression data of the top 2000 most variable features (selection.method = ‘vst’) were scaled, and PCA dimensionality reduction was subsequently performed with npcs = 30 in Seurat 4.3. To integrate expression data from different samples, reciprocal PCA (RPCA) was employed with a configuration of k.anchor = 20, applied separately to blood and tissue samples. Neighborhood graph and uniform manifold approximation and projection (UMAP) and clustering were then performed on the integrated assay in Seurat. Major cell types were annotated based on endometrial scRNA-seq datasets during menstrual cycle and pregnancy (*11, 20, 21*) or known human PBMC markers in the Azimuth database (https://azimuth.hubmapconsortium.org/references/). After clusters of low-quality cells were discarded, 43,037 in blood and 28,876 cells in tissue—each exhibiting over 200 detected genes, below 20% mitochondrial content, and successful donor/recipient assignment—were retained for further analysis. The FindAllMarkers function was utilized to detect marker genes for each annotated cluster with the parameter only.pos = T and a minimal logFC cutoff of 0.5, and the expression of top markers were visualized by ComplexHeatmap 2.16.0. Within major immune cell types, a re-clustering procedure was performed, followed by manual annotation to discern immune subtypes. The visualization of annotated cell clusters and the expression of marker genes on UMAPs were executed utilizing dittoSeq 1.12.2 and Nebulosa 1.10.0, respectively.

### Functional validation of reconstructed immune cells using scRNA-seq data

Enrichment of gene sets, including human TRM signatures (shared DEGs across jejunum, lung, and skin) (*37*), trNK signatures (tissue-specific CD56^bright^ vs CD56^dim^ DEGs in lung or lung lymph node) (*38*), or NK/macrophage signatures derived from Garcia-Alonso et al. and Wang et al., in our data was conducted by GSEA in clusterProfiler and visualized by enrichplot. To draw comparisons in cytokine signaling between reconstructed immune cells and normal immune cells in Garcia-Alonso et al. and Wang et al., gene counts were converted to transcripts per million (TPM) and subjected to a log2 transformation, with subsequent mean centralization conducted across cells. The data were then imported to CytoSig v0.0.3 for analysis, utilizing the parameter - s 2 to incorporate expanded signatures. The obtained activity *Z*-scores were normalized to a 0-1 range across cells to ensure equitable comparisons across datasets, and *P*-values were calculated by comparing Z-scores in tissue-resident cells with those in their circulating counterparts using Student’s *t*-tests. To evaluate the regulatory network activities within UTX/HSCT datasets in comparison to other two datasets, gene regulatory networks in each dataset were identified using the SCENIC package. First, co-expression modules were detected via the ‘pyscenic grn’ command, followed by a pruning of target genes dependent on the presence or absence of specific transcription factor motifs using the ‘pyscenic ctx’ command. Subsequently, the ‘pyscenic aucell’ command was employed to derive Area Under the Curve (AUC) values which measure the enrichment transcription factors’ target genes within expressed genes per regulon per cell, and AUC values were normalized to a range of 0-1 across cells. Differences in regulon activities between tissue-resident and circulating NK cells were assessed through Wilcoxon tests. Finally, to compare cell-cell interactions between our and other datasets, the counts per million (CPM) normalized expression data along with cell type annotation from each dataset were imported to CellphoneDB v4.1.0 for statistical analysis under default settings. The resulting mean values of average ligand and receptor expression per interaction were normalized to a 0-1 range for fair comparisons across datasets. The ktplots 1.2.5 package was utilized to plot *P*-values and normalized mean values, with some custom modifications.

### Statistical analysis

Tests for statistically significant differences were performed for flow cytometric data in Graphpad Prism v10 and for RNA-sequencing data in R. Flow cytometric data was analysed for two-group comparisons with either Mann-Whitney or student t-test depending on verdict from normality test. For multi-group comparisons ANOVA followed by Tukey’s test for multiple comparisons was applied.

### Graphics

Selected schematic figures were either completely or in parts created with BioRender.com.

## Acknowledgements

The authors acknowledge the Bioinformatics and Expression Analysis core facility (BEA), which is supported by the board of research at the Karolinska Institutet and the research committee at the Karolinska University hospital. The authors acknowledge support from the National Genomics Infrastructure in Stockholm funded by Science for Life Laboratory, the Knut and Alice Wallenberg Foundation and the Swedish Research Council, and SNIC/Uppsala Multidisciplinary Center for Advanced Computational Science for assistance with massively parallel sequencing and access to the UPPMAX computational infrastructure. The computations and data handling were enabled by resources provided by the Swedish National Infrastructure for Computing (SNIC) at UPPMAX partially funded by the Swedish Research Council through grant agreement no. 2018-05973. Selected schematic figures were either completely or in parts created with BioRender.com. The authors thank Maria Fursäter and Dr. Aino Fianu Jonasson for help with recruiting study participants.

## Author Contributions

NKB, MAI, and BS planned the study and wrote the manuscript. NKB, BS, and MAI performed experiments and analyzed data, DS and CZ analyzed data, YCG, AB, MS, NM, HK, AFR, SG, and MB contributed to patient recruitment and sample collection. All authors read and gave critical input on the final version of the manuscript.

---

**Table S1.**
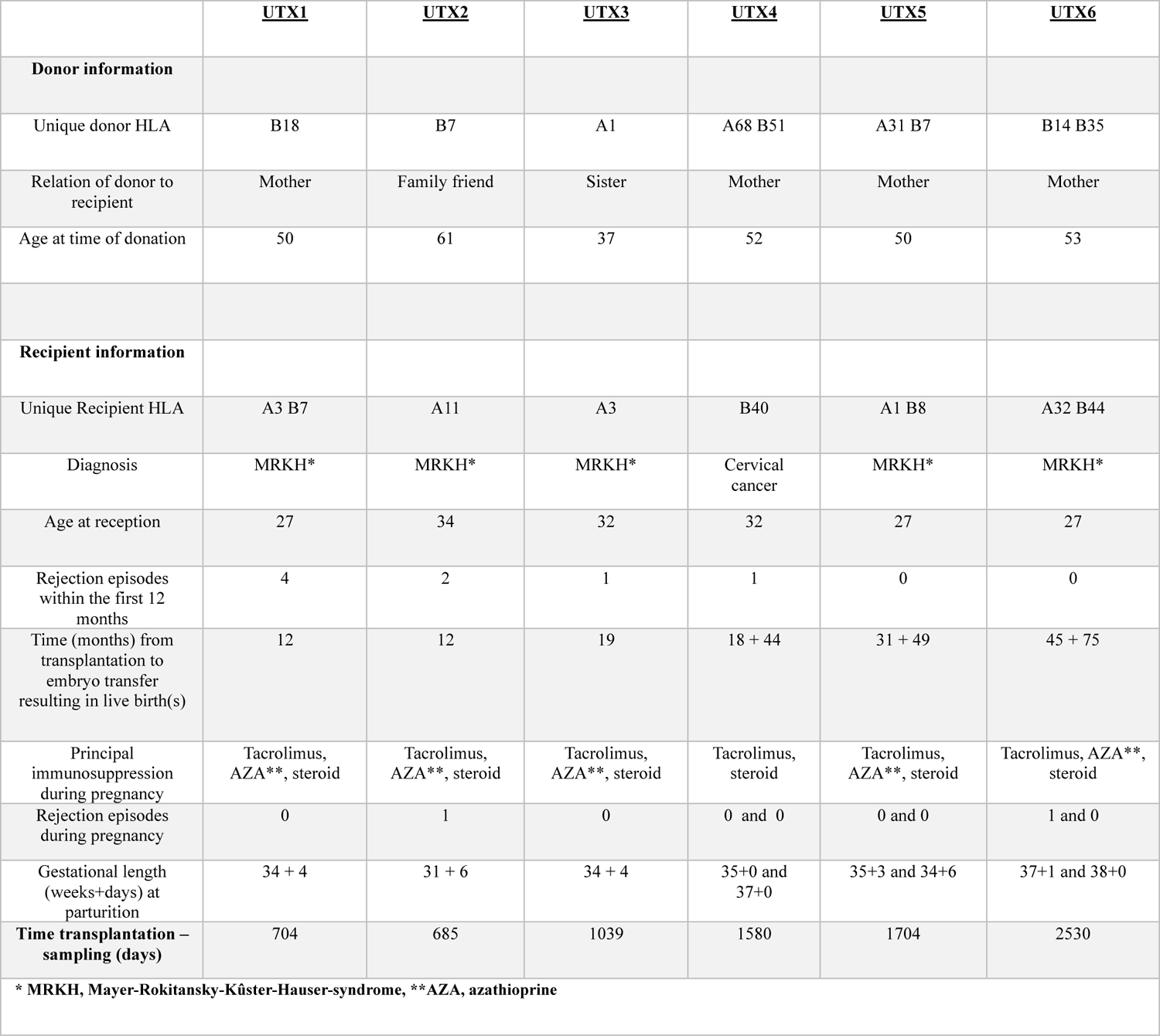
Clinical information of UTX patients.

**Table S2.** Clinical information of HSCT patients.

|  |  |
| --- | --- |
| **Age, years** | 28 (23-47) |
| **Time btw transplantation and menstruation, years** | 0,25- 7 |
| **Time btw transplantation and sampling, years** | 4-11 |
| **Type of transplantation** | allogeneic SCT |
| **Pregnancy y/n** | 3/1 |
| **Type of condition regimen (Reduced intensity conditioning/ Myeloablative)** | 2/2 |
| **Cause for transplantation** | Aplastic anemia, 2x MDS, CML |
| **Donor (Matched unrelated donor / sibling)** | 3/1 |
| **Source of cells (Bone marrow/ peripheral blood stem cells)** | 1/3 |

**Table S3.** Reagents.

| Antigen | Clone | Fluorochrome | Company | Catalogue nr |
| --- | --- | --- | --- | --- |
| anti-IgM | R6-60.2 | BV650 | BD Bioscience | 564027 |
| BDCA-2 | 201A | BV421 | Biolegend | 354211 |
| CCR7 | G043H7 | BV421 | Biolegend | 353208 |
| CD103 | Ber-ACT8 | BB660 | BD Biosciences | 624295 |
| CD103 | Ber-ACT8 | BUV395 | BD Bioscience | 564346 |
| CD11c | 3.9 | PE-Dazzle594 | Biolegend | 301642 |
| CD123 | 6H6 | BV510 | Biolegend | 306022 |
| CD127 | A019D5 | BV711 | Biolegend | 351327 |
| CD138 | MI15 | BV421 | Biolegend | 356516 |
| CD14 | M5E2 | V500 | BD Biosciences | 561391 |
| CD14 | 61D3 | PE-Cy5 | eBioscience | 15-0149-42 |
| CD14 | MφP9 | PE-Cf594 | BD Bioscience | 562334 |
| CD15 | HI98 | V500 | BD Horizon | 561585 |
| CD16 | 3J8 | BUV496 | BD Biosciences | 612944 |
| CD16 | 3G8 | BB755 | BD Biosciences | 624391 |
| CD16 | 3G6 | BV786 | Biolegend | 302046 |
| CD16 | 3J8 | BUV496 | BD Biosciences | 612944 |
| CD16 | 3G8 | BV605 | BD Biosciences | 563172 |
| CD163 | GHI/61 | AF647 | BD Biosciences | 562669 |
| CD18 | REA1112 | PE-Vio615 | Miltenyi | 130-119-090 |
| CD19 | BV510 | SJ25C1 | BD Biosciences | 562947 |
| CD19 | HIB19 | PE-Cy5 | Biolegend | 302210 |
| CD19 | SJ25C1 | BV786 | BD Biosciences | 563325 |
| CD206 | DCN228 | PE-VIO770 | Miltenyi | 130-130-366 |
| CD209 | REA617 | APC-Vio770 | Miltenyi | 130-120-729 |
| CD25 | 2A3 | BUV395 | BD Biosciences | 564034 |
| CD27 | 2M-T271 | BUV496 | BD Biosciences | 741145 |
| CD3 | SK7 | BV750 | Biolegend | 344845 |
| CD3 | UCHT1 | PC5 | Beckman Coulter | 6607010 |
| CD3 | UCHT1 | PE-Cy5 | BioLegend | 300410 |
| CD38 | HIT2 | BUV661 | BD Biosciences | 612969 |
| CD39 | TU66 | BB755 | BD Biosciences | 624391 |
| CD39 | TU66 | BV711 | BD Bioscience | 563680 |
| CD4 | SK3 | BUV615 | BD Biosciences | 612987 |
| CD4 | OKT4 | PE-Cy5 | Biolegend | 317412 |
| CD45 | HI30 | BUV805 | BD Biosciences | 612891 |
| CD45 | HI30 | AF700 | Biolegend | 304024 |
| CD45 | HI30 | BV785 | Biolegend | 304048 |
| CD46RA | HI100 | BV785 | Biolegend | 304140 |
| CD49a | SR84 | BUV615 | BD Biosciences | 624297 |
| CD49a | TS2/7 | APC | Biolegend | 328310 |
| CD49e | IIA1 | BUV563 | BD Biosciences | 741383 |
| CD56 | NCAM16.2 | BUV737 | BD Biosciences | 564447 |
| CD56 | N901 | ECD | Beckman Coulter | A82943 |
| CD56 | 5.1H11 | BV785 | Biolegend | 362550 |
| CD57 | TB03 | APC-Vio770 | Miltenyi | 130-116-503 |
| CD57 | REA769 | APC-Cy7/H7 | Miltenyi | 130-111-813 |
| CD57 | TB01 | Purified | Thermo Fisher | 16-0577-85 |
| CD69 | FN50 | Biotin | Biolegend | 310924 |
| CD69 | FN50 | BUV737 | BD Bioscience | 612817 |
| CD8 | RPA-T8 | BV570 | Biolegend | 301037 |
| CD8 | RPA-T8 | BB700 | BD Optibuild | 566452 |
| CD8 | 3B5 | QD605 | Thermo Fisher | Q10009 |
| CD9 | REA1071 | APC-Vio770 | Miltenyi | 130-118-813 |
| CD9 | M-L13 | BUV615 | BD Biosciences | 624297 |
| DNAM-1 | 11A8 | PE-Cy7 | Biolegend | 338306 |
| EB6 | EB6B | PE-Cy5.5 | Beckman<br>Coulter | a66898 |
| HLA-A1, 36 | NA | Biotin | OneLambda | BIH0331 |
| HLA-A2 | BB7.2 | FITC | Biolegend | 343304 |
| HLA-A2, 28 | REA142 | Biotin | Miltenyi | 130-099-594 |
| HLA-A3 | REA950 | FITC | Miltenyi | 130-115-739 |
| HLA-B7, 27 | REA176 | FITC Viobright | Miltenyi | 130-120-234 |
| HLA-B8 | REA145 | Biotin | Miltenyi | 130-099-588 |
| HLA-DR | L243 | BV785 | Biolegend | 307642 |
| IgD | IA6-2 | BUV395 | BD Horizon | 563813 |
| KIR2DL1/S1 | EB6B | PE-Cy5.5 | Beckman<br>Coulter | A66898 |
| KIR2DL1/S1 | EB6B | PE-Cy7 | Beckman<br>Coulter | A66899 |
| KIR2DL2/L3/S2 | GL183 | PE-Cy5.5 | Beckman<br>Coulter | A66900 |
| KIR2DL3 | 180701 | FITC | RnD Systems | FAB2014F |
| KIR2DS4 | 179315 | In house<br>biotinylated | RnD Systems | MAB1847 |
| KIR3DL1 | Dx9 | BV421 | Biolegend | 312714 |
| LILRB1 | 292305 | APC | RnD Systems | FAB20171A |
| NKG2A | REA110 | PE | Miltenyi | 130-113-566 |
| NKG2A | REA110 | APC | Miltenyi | 130-113-563 |
| Streptavidin | NA | BV650 | BD Biosciences | 563855 |
| Streptavidin | NA | QD585 | Life<br>Technologies | Q10111MP |
| Streptavidin | NA | BV650 | BD Bioscience | 563855 |
| TCRgd | IMMU510 | PE-Cy5.5 | Beckman<br>Coulter | A99021 |
| TCRVa7.2 | 3C10 | PE | Biolegend | 351706 |

**Fig. S1.**
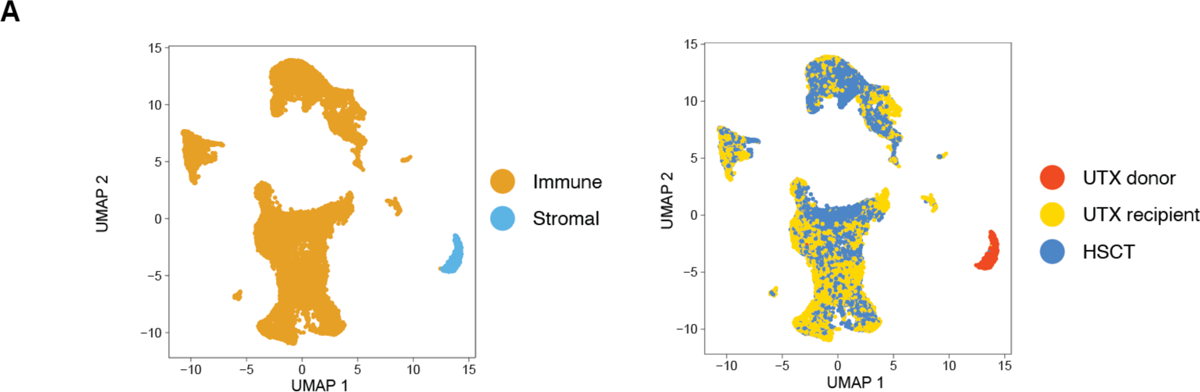
Origin of stromal cells in scRNAseq data of UTX and HSCT patients. Displayed as UMAP projection are the annotation to immune or stromal cells (left hand side) or the allocation as either donor or recipient derived cells in UTX or HSCT (right hand side).

**Fig. S2.**
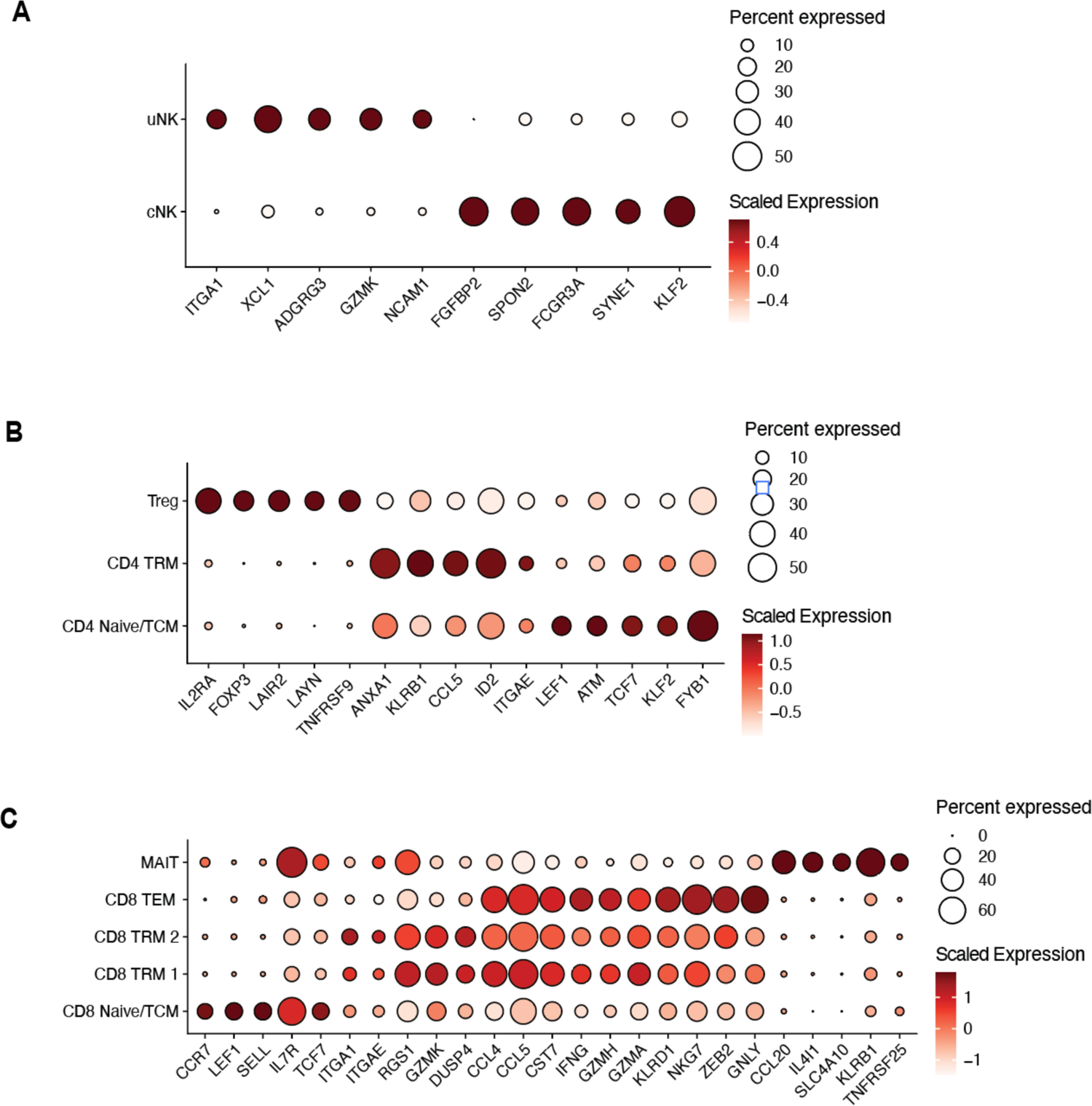
Identification of major immune cell subsets in uterine samples from UTX and HSCT patients. (A-C) Display of differentially expressed genes based on scRNAseq of UTX and HSCT samples identifying the indicated subsets in NK cells (A), CD4 T cells (B) and CD8 T cells (C).

**Fig. S3.**
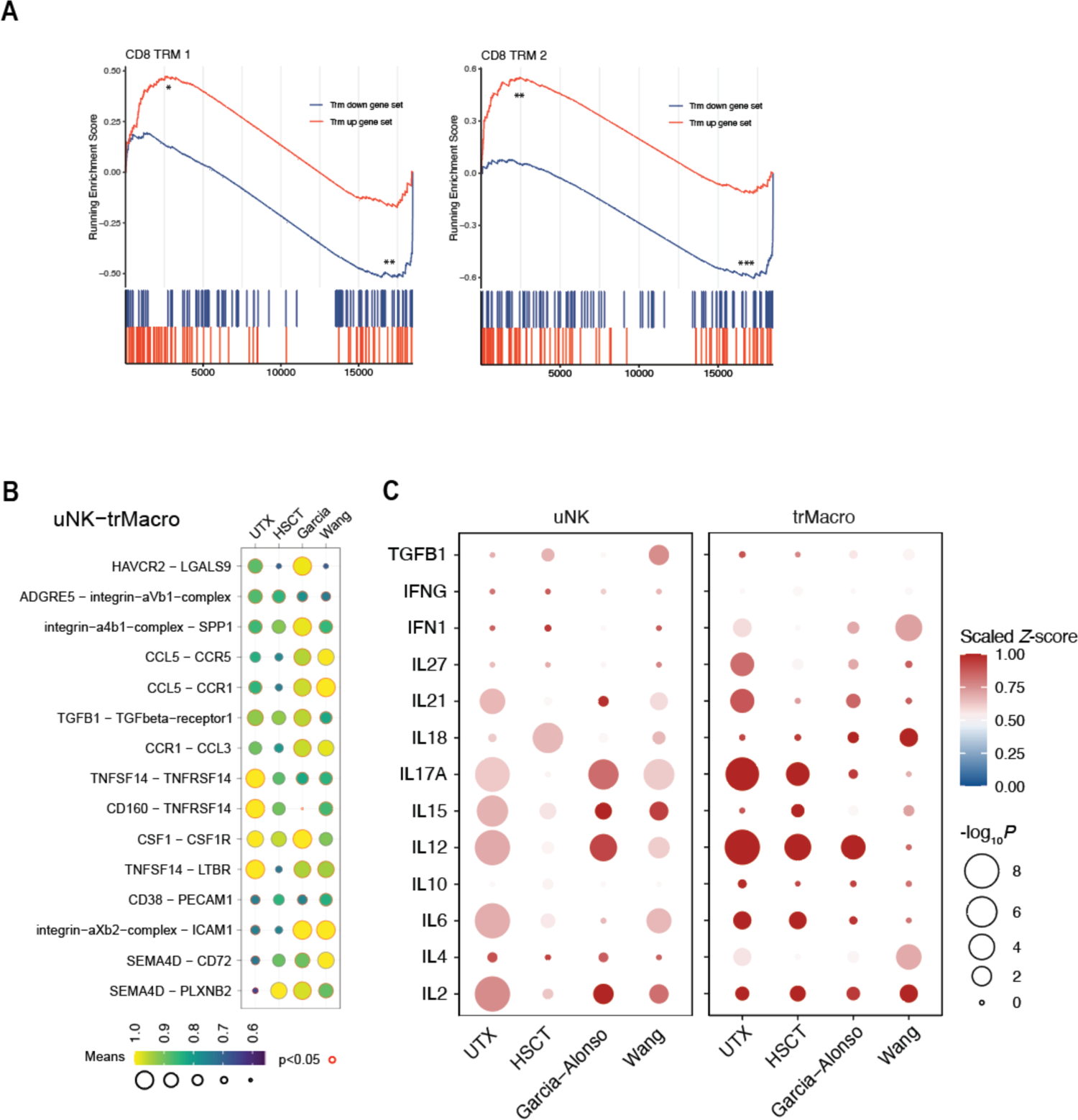
Tissue residency profiles and cell-cell communication after transplantation. (A) Gene set enrichment analysis (GSEA) for tissue residency profiles of CD8 TRM subsets based on scRNAseq data. (B and C) Analysis of integrated scRNAseq data, comprising uterus transplantation (UTX), hematopoietic stem cell transplantation (HSCT) and control datasets (*Garcia-Alonso et al*., 2021, *Wang et al*., 2020). (**B**) Display of top 15 cellular interactions determined with Cellphone-DB for uterine NK (uNK) cells and tissue-resident macrophages (trMacro). (**C**) Cytokine-related pathways determined with Cytosig in uNK and trMacro in the indicated datasets.

**Fig. S4.**
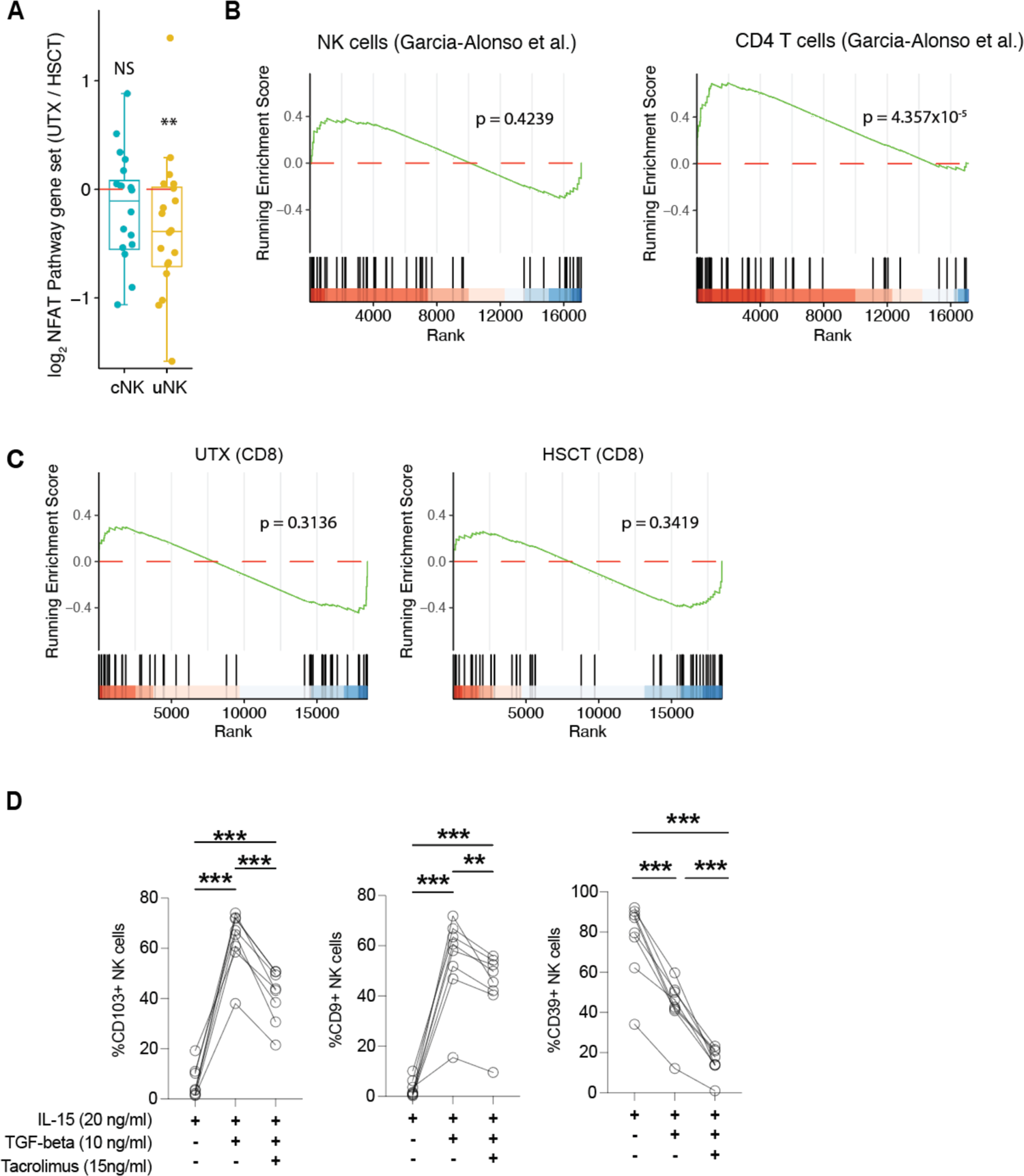
NFAT signaling in NK and T cells in the uterus. (A) Enrichment of the NFAT pathway gene set in UTX/HSCT comparing uterine and circulating NK cells (uNK and cNK, respectively). (B and C) GSEA of NFAT signaling pathway in the indicated subsets based on scRNAseq data from Garcia-Alonso et al (B) or in CD8 TRM cells in either UTX or HSCT patients (C). (D) Expression of the indicated markers after in vitro stimulation with combinations of IL-15, TGF-beta and Tacrolimus in total NK cells. For comparison of three groups ANOVA followed by Tukey’s test for correction of multiple comparisons was used, * indicates p<0.05, ** p<0.01 and *** p<0.001.

**Fig. S5.**
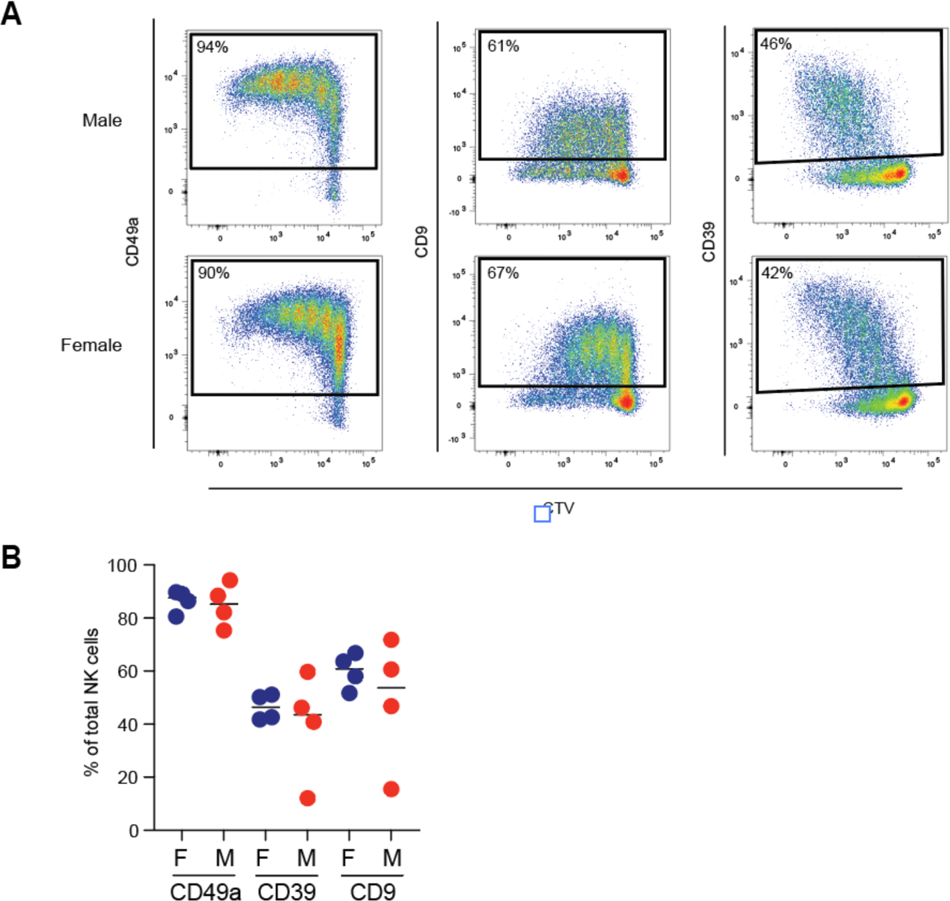
Effect of genetic sex on formation of uterine NK cell phenotype. (A-B) Representative flow cytometry staining (A) and summary data (B) of indicated marker expression on NK cells after in vitro stimulation with IL-15 and TGF-beta, stratified for genetic sex of NK cells.

